# HDAC6 inhibition partially alleviates mitochondrial trafficking defects and restores motor function in human motor neuron and zebrafish models of Charcot-Marie-Tooth Disease Type 2A

**DOI:** 10.1101/2022.07.05.498819

**Authors:** Larissa Butler, Kathryn I. Adamson, Stuart L. Johnson, Lydia H. Jestice, Christopher J. Price, Dylan Stavish, Niedharsan Pooranachandran, Jarema J. Malicki, Anestis Tsakiridis, Andrew J. Grierson, Ivana Barbaric

## Abstract

Charcot Marie Tooth Disease (CMT) is a group of inherited progressive conditions affecting distal motor and sensory neurons, leading to muscle weakness, pain and loss of sensation in limbs. There are currently no treatments for this debilitating disease. To investigate disease mechanisms and facilitate treatment discovery, here we developed an *in vitro* model for CMT type 2A by introducing the patient-specific *MFN2*^R94Q^ mutation into human embryonic stem cells (hESCs). Isogenic mutant and wild-type hESCs differentiated to spinal motor neurons with similar efficiency and gave rise to functional motor neurons *in vitro*. However, MFN2^R94Q/+^ spinal motor neurons displayed impaired mitochondrial trafficking, resulting in reduced numbers of mitochondria in distal parts of axons. Importantly, we showed that mitochondrial trafficking defects can be alleviated by treatment with an HDAC6 inhibitor. Chemical and genetic inhibition of HDAC6 also significantly rescued the motor phenotype in a zebrafish CMT2A model. Taken together, our study reveals a mutation-specific insight into CMT2A disease mechanism and confirms HDAC6 as a promising target for further therapeutic development.

## Introduction

Charcot Marie Tooth Disease (CMT) is a group of inherited progressive conditions affecting spinal motor and sensory nerves, resulting in muscle weakness, pain and loss of sensation in limbs of CMT patients (Reilly et al., 2011). Traditionally, CMT is classified into several different types, with the onset of some of the most severe clinical manifestations occurring within the subtype CMT2A (Feely et al., 2011; Reilly et al., 2011). CMT2A patients typically become wheelchair-reliant by the age of 20 and often exhibit optical atrophy and/or hearing loss (Feely et al., 2011). CMT2A is caused by mutations in Mitofusin 2 (*MFN2*), which encodes a GTPase protein located on the outer mitochondrial transmembrane (Zuchner et al., 2004). MFN2 has several known roles, including mediating the outer membrane fusion of mitochondria (Chandhok et al., 2018). Over 100 different mutations have been mapped to MFN2 in CMT2A patients (Stuppia et al., 2015; Verhoeven et al., 2006). A particularly frequent and phenotypically severe form of the disease is caused by a missense mutation leading to the substitution of arginine with either glutamine or tryptophan at the residue 94 of MFN2 (denoted as R94Q and R94W, respectively) (Feely et al., 2011; Verhoeven et al., 2006). Despite significant progress in the identification of MFN2 mutations in CMT2A patients, how such mutations impact MFN2 function to give rise to the disease phenotype remains poorly understood.

In CMT2A, progressive degeneration of spinal motor and sensory neurons affects the lower extremities earlier and more severely than the upper extremities, consistent with axonal length playing a role in the CMT2A disease mechanism. An attractive hypothesis to explain the nerve length-dependent manifestation of CMT2A focuses on aberrant mitochondrial trafficking through axons (Dorn, 2020; Prior et al., 2017). A seminal study in cultured rat DRG neurons expressing a mutant form of MFN2 demonstrated an abnormal clustering of mitochondria in neuronal cell bodies and proximal axons as well as the reduced mitochondrial mobility in mutant cells (Baloh et al., 2007). Subsequent studies in other *in vivo* and *in vitro* models of CMT2A, including zebrafish and mouse (Chapman et al., 2013; Rocha et al., 2018), supported the observation of impaired mitochondrial trafficking caused by *MFN2* mutations. Consequently, rescuing axonal trafficking of mitochondria has been highlighted as a promising therapeutic strategy for the treatment of CMT2A (Rossaert and Van Den Bosch, 2020). An HDAC6 inhibitor, tubastatin A, effectively alleviated motor and sensory defects in a mouse model of CMT type 2F (CMT2F) by increasing α-tubulin acetylation in peripheral nerves of mutant mice (d’Ydewalle et al., 2011). HDAC6 inhibition also reversed mitochondrial trafficking defects in a CMT type 2D (CMT2D) mouse model (Benoy et al., 2018). Moreover, HDAC6 inhibitors proved effective in alleviating mitochondrial transport in models of other neurodegenerative diseases whose pathology is also linked to mitochondrial trafficking, such as amyotrophic lateral sclerosis (Fazal et al., 2021; Guo et al., 2017). Together, these observations suggest that the HDAC6 inhibition may provide a much-needed therapeutic option for neurodegenerative diseases underpinned by mitochondrial transport defects. Nonetheless, the impact of HDAC6 inhibition is yet to be tested in models of the most severe CMT2 type, CMT2A.

The selective vulnerability of spinal motor neurons to CMT2A necessitates the use of disease-relevant cell types for the investigation of MFN2 disease-associated dysfunction and for testing approaches for alleviating the disease phenotype. While human spinal motor neurons are experimentally inaccessible, the advent of human pluripotent stem cell (hPSC) technology has offered a means to derive them *in vitro*. HPSCs, which encompass both human embryonic stem cells (hESCs) derived from early blastocysts (Thomson et al., 1998) and induced pluripotent stem cells (hiPSC) reprogrammed from somatic cells (Takahashi et al., 2007), have the ability to differentiate to any cell type (Takahashi et al., 2007; Thomson et al., 1998). Here, we introduced the patient-specific mutation *MFN2*^R94Q/+^ into hESCs to obtain isogenic mutant and wild-type cells and we established a robust protocol for differentiation of hPSCs to disease-relevant spinal motor neurons. We show that impaired mitochondrial trafficking and distally altered mitochondrial morphology are the key features of MFN2^R94Q/+^ motor neurons and that these trafficking defects can be reversed by chemically inhibiting HDAC6. Furthermore, to demonstrate the relevance of these findings *in vivo*, we show that chemical or genetic inhibition of HDAC6 in a zebrafish MFN2 mutant (Chapman et al., 2013) significantly rescued their aberrant swimming phenotype, thus confirming HDAC6 as a promising therapeutic target in CMT2A.

## Results

### Generation of isogenic MFN2^R94Q/+^ and MFN2^+/+^ hESCs

We set out to generate a human cell-based model of CMT2A by introducing one of the most common and phenotypically severe CMT2A-causing mutations, MFN2^R94Q/+^, into a hESC line MShef11 (Thompson et al., 2020). To this end, we transfected euploid early-passage hESCs with a Cas9-guide RNA duplex along with a repair template designed to introduce the G281A mutation into *MFN2* **(Figure 1A, B)**. To generate clonal lines, bulk transfected cells were single-cell sorted and expanded **(Figure 1B)**. The resulting hESC colonies were subsequently screened for successful editing by Sanger sequencing of the target region. Following the screening, two heterozygous edited clones (from herein termed MFN2^R94Q/+^ Het1 and MFN2^R94Q/+^ Het2) were identified and expanded for in-depth analyses **(Figure 1B,C)**. We further validated successful heterozygous MFN2^R94Q/+^ editing by re-cloning and sequencing sub-clones of the two mutant lines **(Supplementary Figure 1A)**. As control cells in phenotypic assays, we used wild-type parental cells (termed MFN2^+/+^ Ctrl1) and an unedited wild-type clone that has undergone CRISPR/Cas9 transfection (termed MFN2^+/+^ Ctrl2) **(Figure 1B,C)**. All of the lines remained karyotypically diploid and did not harbour any predicted off-target mutations arising during CRISPR/Cas9 mutagenesis **(Supplementary Figure 1B)**. These lines also retained their undifferentiated hESC phenotype post-editing, as evidenced by colony morphology **(Supplementary Figure 1C)** and expression of key pluripotency-associated markers **(Figure 1D, E)**. Overall, the CRISPR/Cas9 editing of hESCs enabled us to generate isogenic lines with and without MFN2^R94Q/+^ mutation for CMT2A disease modelling.

**Figure 1.**
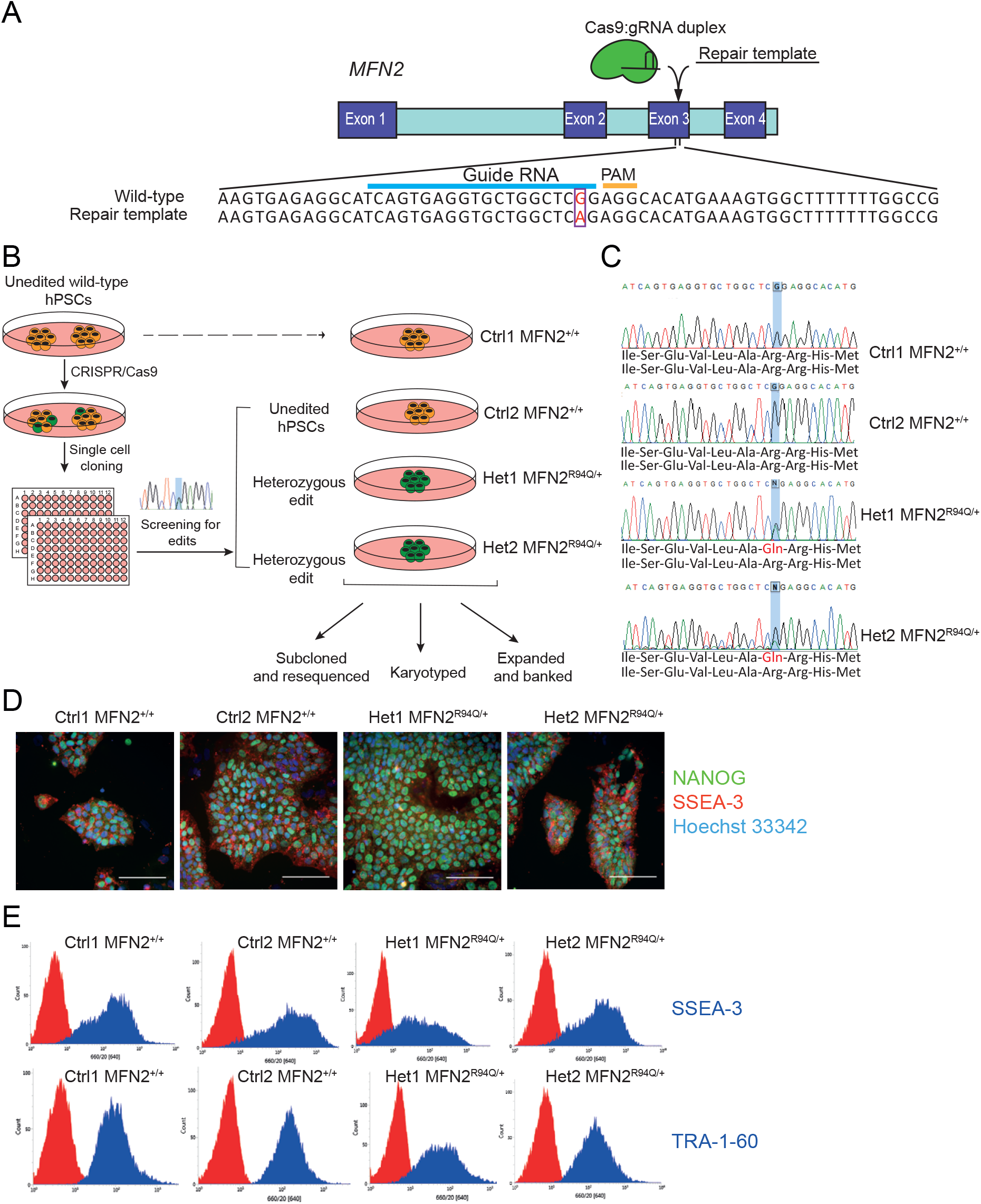
Generation of isogenic MFN2^R94Q/+^ and MFN2^+/+^ hESC clones using CRISPR/Cas9. **A)** Schematic representation of the gene editing strategy. Guide RNA and PAM sequences are shown alongside the wild-type genomic sequence of *MFN2* Exon 3 containing the intended nucleotide for editing. Intended nucleotide for editing is boxed and labelled in red in wild-type and repair template sequences, G to A, respectively. **B)** Schematic depicting workflow for the genome editing and cloning of edited and control sublines. **C)** DNA sequence analysis verified successful genome editing, with two peaks in the MFN2^R94Q/+^ clones (G and A, highlighted in the sequence) indicating heterozygous edit. Predicted amino acid sequence is shown below the sequence traces. **D)** Representative images of MFN2^R94Q/+^ and MFN2^+/+^ hPSC stained for NANOG (green) and SSEA3 (red) markers of undifferentiated state. Nuclei are counterstained with Hoechst 33342. Scale bar: 100μm. **E)** Representative flow cytometry plots of SSEA3 and TRA-1-60 markers of undifferentiated hESC state (blue histogram) against an isotype control (red histogram).

### MFN2^R94Q/+^ hESCs efficiently differentiate into functional spinal motor neurons

To model CMT2A *in vitro*, we aimed to produce spinal motor neurons from MFN2^R94Q/+^ hESCs. During development, limb-innervating spinal motor neurons arise in the brachial and lumbar-level spinal cord (Stifani, 2014). They are characterised by high expression of FOXP1 (Dasen et al., 2008), in addition to the expression of motor neuron markers (e.g. choline acetyltransferase (ChAT), ISLET-1/2, OLIG2, and HB9 (MNX1)) (Mizuguchi et al., 2001; Rayon et al., 2021). OLIG2 is a motor neuron progenitor marker, which induces the expression of post-mitotic marker HB9 (Lee and Pfaff, 2003). HB9 expression is downregulated in mature neurons targeting the dorsal side of the limb, whereas HB9 and LHX1 are expressed in ventrally targeting populations (Rousso et al., 2008). Depending on their axial level, limb-innervating motor neurons express different HOX paralogous group (PG) genes. For example, brachial motor neurons express HOX PG 6 members, with HOX PG 5 expressed rostrally and HOX PG 8 caudally (Dasen and Jessell, 2009), whereas lumbar motor neurons express HOX PG 10 (Dasen and Jessell, 2009).

To develop an efficient protocol for limb-innervating spinal motor neuron differentiation, we modified previously published protocols for neural differentiation from hESCs relying on the induction of an intermediate populations resembling neuromesodermal-potent axial progenitors (NMPs), the bona fide precursors of the embryonic spinal cord (Gouti et al., 2014; Guo et al., 2017; Maury et al., 2015). Our optimisation involved extending the timing of the dual SMAD inhibition to facilitate the induction of neural identity (Chambers et al., 2009). Additionally, we extended and increased the concentration of CHIR99021 (WNT agonist) in combination with FGF for the first four days of the protocol. The combination of WNT/FGF is key for the induction of NMPs and posterior *HOX* genes (Gouti et al., 2017). The final alteration made to the protocol was to increase the concentration of all-*trans* retinoic acid, as this is a key molecule in the neuralisation of NMPs and additionally has roles in *HOX* gene regulation (Gouti et al., 2017; Simeone et al., 1990) **(Figure 2A)**. At days 2 and 4 after the initiation of differentiation, we noted significant upregulation of *CDX2, TBXT* and *NKX1*.*2*, as well as an upregulation of *HOX* transcripts, indicating the acquisition of neuromesodermal progenitor/ early spinal cord identity (**Figure 2B)**. After 16-day culture of MFN2^+/+^ Ctrl1 in the modified differentiation conditions, the majority of the cells were positive for ChAT, ISLET-1/2, OLIG2 and HB9 (**Figure 2C**), consistent with motor neuronal induction. Further differentiation in the modified differentiation conditions resulted in robust induction of motor neuron markers ChAT (81 ±13%), ISLET-1/2 (76 ±10%) and HB9 (92 ±5%) at day 33 (**Figure 2D,E)**. The majority of obtained neurons expressed FOXP1 (77 ±20%) (**Figure 2D,E**), consistent with the limb-innervating motor neuron fate (Rousso et al., 2008; Stifani, 2014), although we also detected presence of *PHOX2B*, a marker of more anterior, cervical-level motor neurons (**Figure 2F)**. Further analysis of the gene expression also confirmed the phenotype consistent with a lateral motor column identity, including high expression of *ISLET-1/2* and *HB9* motor neuron markers (**Figure 2F**). Finally, we detected a significant increase in expression of *HOXA4, HOXC6* and *HOXC8* at both day 13 and day 33 of differentiation compared to undifferentiated hESCs (**Figure 2G**). Taken together, this protocol enabled us to differentiate hESCs to brachial-level spinal motor neurons.

**Figure 2.**
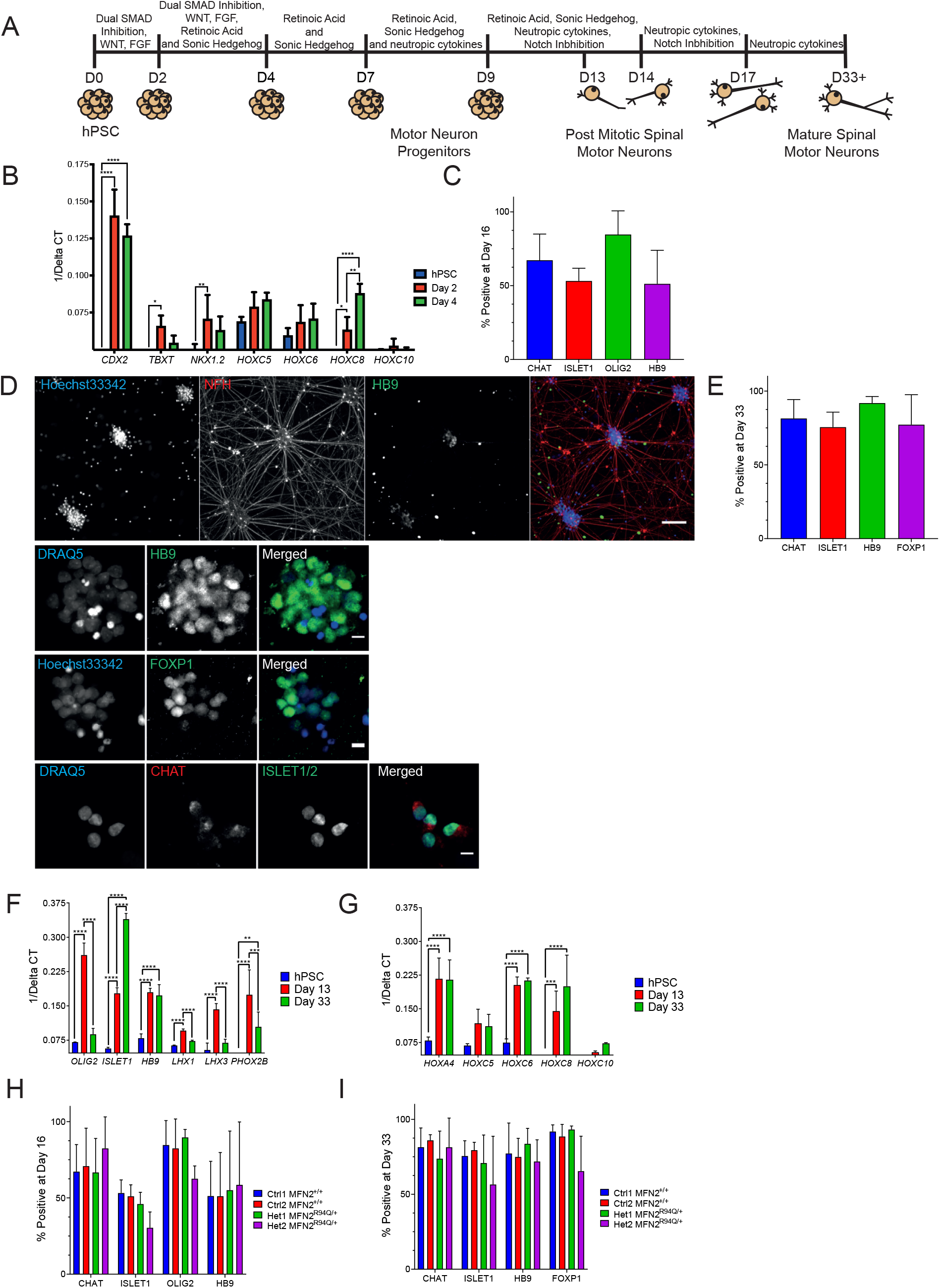
Differentiation of hESCs to spinal motor neurons. **A)** Schematic depicting optimised differentiation protocol of hESC to spinal motor neurons. **B)** Gene expression profiling by qPCR at day 0 (hESCs), day 2 and day 4 after the start of the differentiation protocol, demonstrating the acquisition of neuromesodermal progenitor fate. *p<0.01, **p<0.01, ***p<0.001, ****p<0.0001, 2-way ANOVA. **C)** Quantification of immunocytochemistry for motor neuron markers at day 16 of differentiation of Ctrl1 MFN2^+/+^ hESC. Data shown are the mean ±SD of three biological repeats. **D)** Quantification of immunocytochemistry for motor neuron markers at day 33 of differentiation of Ctrl1 MFN2^+/+^ hESC. **E)** Immunocytochemistry for motor neuron markers at day 33 of differentiation of Ctrl1 MFN2^+/+^ hESC. Nuclei are counterstained with DRAQ5 or Hoechst 33342, as indicated. Scale bar: 100μm for the top-most panel and 10 μm for the remaining panels. **F)** Gene expression profiling by qPCR for Ctrl1 MFN2^+/+^ differentiated cells, demonstrating further expression of relevant motor neuron markers (*OLIG2, ISLET1, HB9*) and columnar markers (*LHX1, LHX3, PHOX2B*); **p<0.01, ***p<0.001, ****p<0.0001, 2-way ANOVA. **G)** Gene expression profiling by qPCR demonstrating further expression of HOX markers from anterior/cranial (*HOXA4*) to posterior/lumbar (*HOXC10*); **p<0.01, ***p<0.001, ****p<0.0001, 2-way ANOVA. **H)** Quantification of immunocytochemistry for motor neuron markers for control and mutant lines at day 16 of differentiation. **I)** Quantification of immunocytochemistry for motor neuron markers for control and mutant lines at day 33 of differentiation.

Next, we ascertained whether the MFN2^R94Q/+^ mutation impacts the differentiation of hESCs to spinal motor neurons. We subjected MFN2^R94Q/+^ Het1 and Het2 hESCs, alongside the MFN2^+/+^ Ctrl1 and Ctrl2 cells, to our optimised differentiation protocol. To compare the differentiation efficiency of MFN2^R94Q/+^ Het1 and Het2 versus MFN2^+/+^ Ctrl1 and MFN2^+/+^ Ctrl2 cells, we quantified the number of cells expressing different motor neuron markers at day 16 (**Figure 2H**) and day 33 (**Figure 2I)** of the differentiation protocol. All of the populations exhibited a similar overall trend in the expression of key markers of motor neuron identity and there were no significant differences in the differentiation efficiency of MFN2^R94Q/+^ Het1 and Het2 cells compared with MFN2^+/+^ Ctrl1 and Ctrl2 hESCs.

Finally, we assessed the functional properties of hESC-derived motor neurons. Motor neurons were found to be electrophysiologically active based on analysis of action potential activity and ionic currents, with no overt differences observed between MFN2^+/+^ and MFN2^R94Q/+^ motor neurons **(Supplementary Figure 2)**. Taken together, this data demonstrates that MFN2^R94Q/+^ mutation does not impact the differentiation efficiency or electrophysiological properties of hESCs-derived spinal motor neurons.

### MFN2^R94Q/+^ impairs mitochondrial trafficking in hESC-derived spinal motor neurons

To directly test the hypothesis that MFN2^R94Q/+^ impacts mitochondrial axonal transport in human spinal motor neurons, we performed time-lapse microscopy of mitochondrial movement within the axons of MFN2^R94Q/+^ and MFN2^+/+^ motor neurons. Motor neurons were co-transfected with a construct encoding a mitochondrial targeting sequence fused to a red fluorescent protein (Mito-DsRed2) and with another construct encoding enhanced green fluorescent protein (eGFP). Only a small proportion of neurons were labelled with both eGFP and Mito-DsRed2, thus enabling us to track mitochondria within individual axons of densely populated motor neuron cultures **(Supplementary Movie S1, S2)**. We considered mitochondria motile if they moved with a speed of over 0.3μm/s, as movements below this threshold could be attributed to actin-mediated transport (De Vos and Sheetz, 2007). Mitochondria in MFN2^R94Q/+^ hESC-derived motor neurons were overall less motile (**Figure 3A,B)** and their transport was reduced in both anterograde (**Figure 3C)** and retrograde (**Figure 3D)** direction compared to mitochondria in wild-type motor neurons.

**Figure 3.**
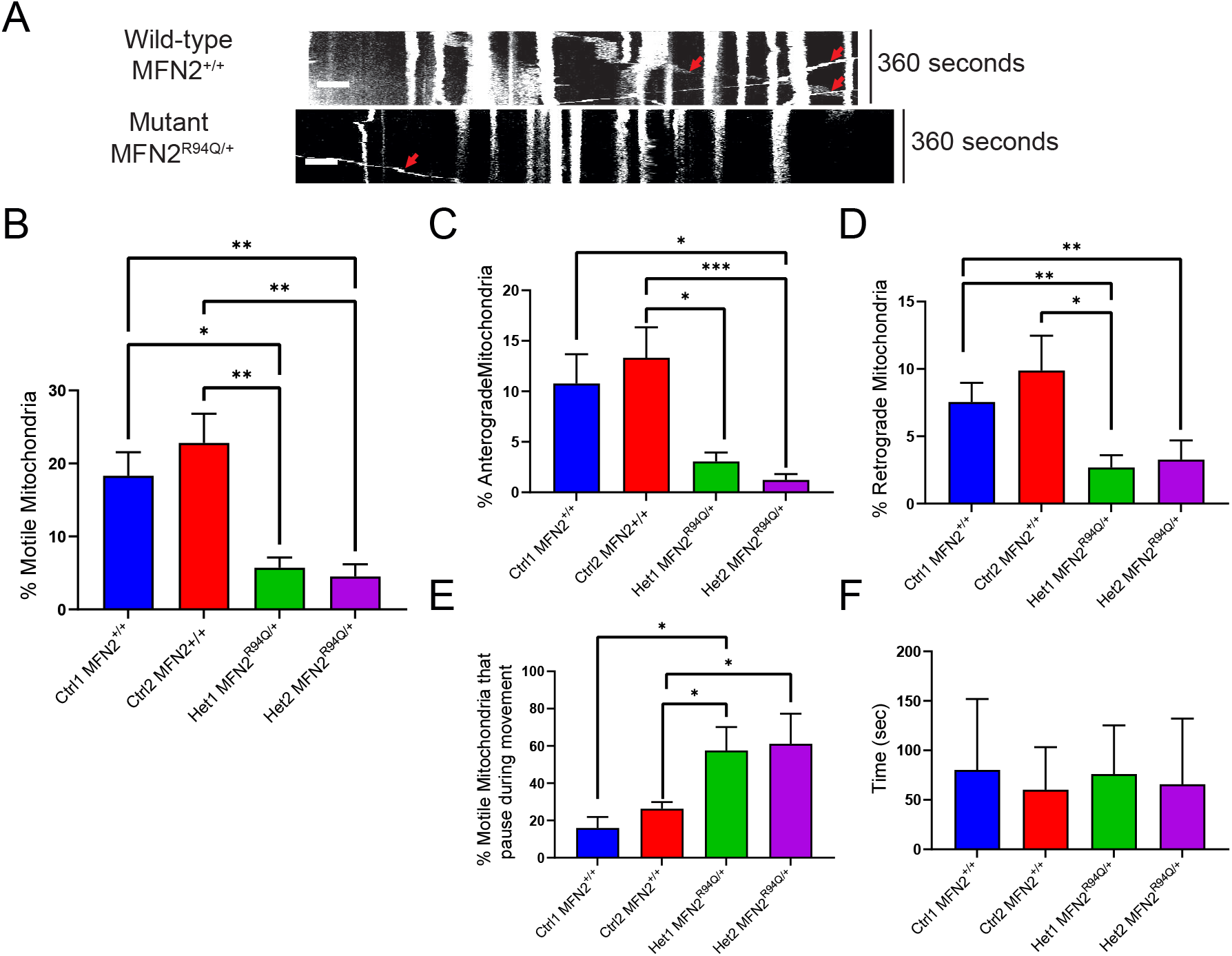
Altered mitochondrial trafficking in MFN2^R94Q/+^ motor neurons. **A)** Representative kymographs of wild-type and mutant neurons. Stationary mitochondria are visible as straight lines. Mitochondrion in motion is depicted as diagonal lines. Arrows indicate several mitochondria in motion. Horizontal scale bar 10 μm, vertical scale 360 seconds. **B)** The percentage of motile mitochondria in control and mutant lines, moving in either anterograde or retrograde direction *p<0.05, **p<0.01, Kruskal-Wallis test. **C)** The percentage of motile mitochondria in control and mutant lines moving in anterograde direction. **D)** The percentage of motile mitochondria in control and mutant lines moving in anterograde direction. **E)** The percentage of mitochondria that pause during movements. **F)** The average time mitochondria spend paused during movements. All data shown are the mean ±SEM. N = 15+ neurons for trafficking data. *p<0.05, **p<0.01, ***p<0.001, Kruskal-Wallis test (B) or Kolmogorov-Smirnov test (C-E).

We next assessed the pausing of mitochondria in wild-type and MFN2^R94Q/+^ hESC-derived motor neurons. Increased pausing can indicate that mitochondria are unable to sustain continued or optimal movement speeds and is therefore associated with mitochondrial transport dysfunction (Misko et al., 2010). In our analysis, mitochondrial pausing was defined as a transient cessation of mitochondrial movement that lasted at least 6s before a mitochondrion restarted movement at a speed higher than 0.3 μm/s. The percentage of motile mitochondria that paused during transport was found to be significantly increased in MFN2^R94Q/+^ hESC-derived motor neurons versus control **(Figure 3E)**, although the average amount of time that mitochondria of mutant cells spent pausing was similar to their wild-type counterparts **(Figure 3F)**. Therefore, our data indicates that mitochondrial transport was more frequently interrupted in the MFN2^R94Q/+^ neurons.

Consistent with the impaired mitochondrial trafficking, we found a decreased number of mitochondria in distal regions of MFN2^R94Q/+^ motor neuron axons in comparison to wild-type counterparts **(Figure 4A,B)**. In addition, the overall spacing between mitochondria was increased in distal regions of MFN2^R94Q/+^ motor neuron axons **(Figure 4C)**, a result in agreement with the observed decrease in mitochondrial numbers. As mitochondrial number and spacing in distal neurons is dependent on mitochondrial transport (Mandal and Drerup, 2019), these results reinforced the notion that *MFN2*^R94Q/+^ mutation impacts the mitochondrial transport along motor neuron axons. Finally, we also noted a significantly reduced aspect ratio of mitochondria in the distal segments of MFN2^R94Q/+^ motor neuron axons **(Figure 4D)**. This phenotype was particularly prominent in the distal part of the axon, indicating that a lower number of mitochondria in distal axons may result in reduced mitochondrial fusion in these axonal regions.

**Figure 4.**
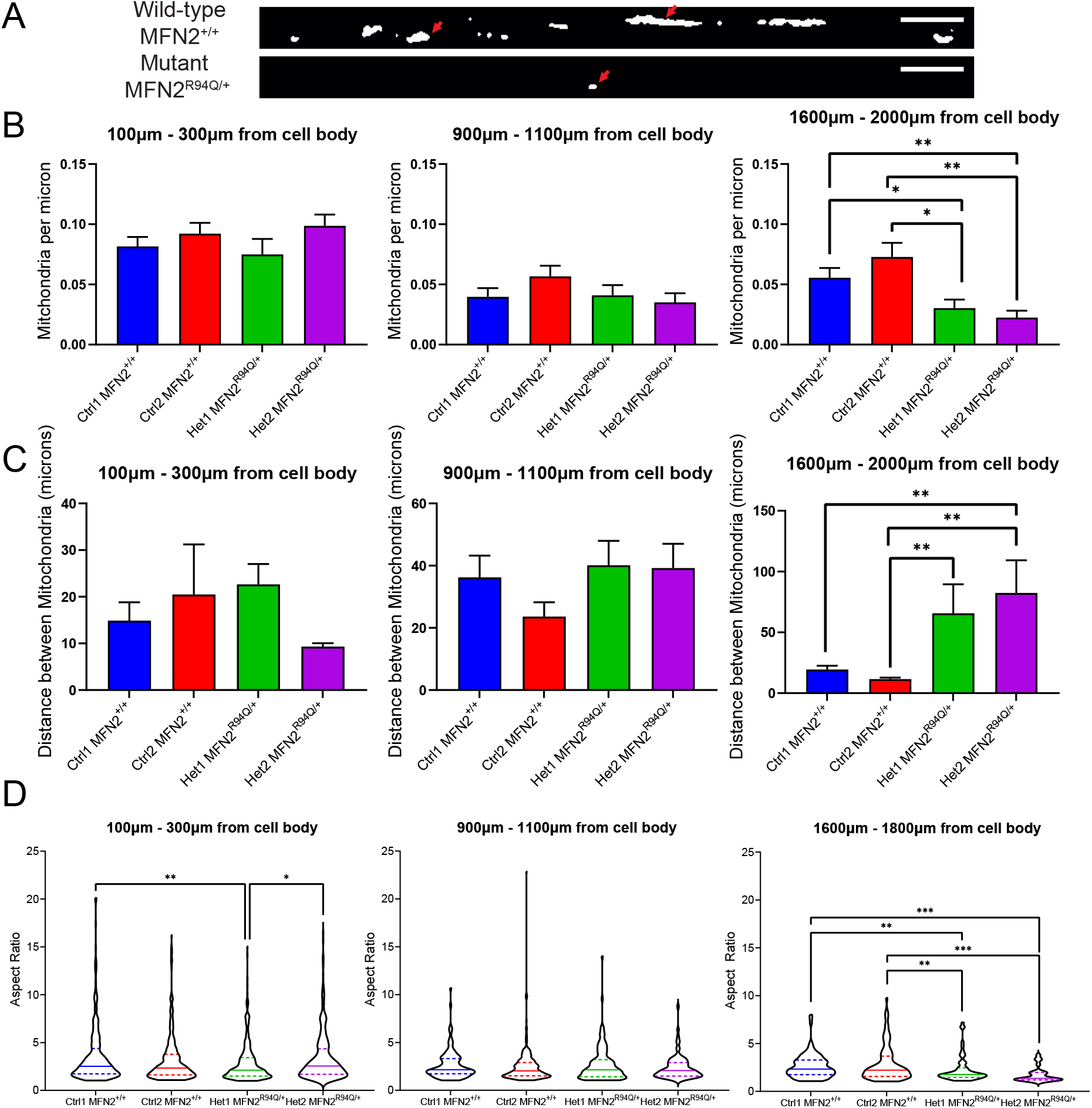
Altered mitochondrial morphology in distal parts of MFN2^R94Q/+^ motor neurons. **A)** Representative images of mitochondria from a distal region of MFN2^+/+^ and MFN2^R94Q/+^ neurons. Mitochondria are seen as white regions in the images. Red arrows point to typical examples of mitochondria seen in each genotype. Note the reduced number and length of mitochondria in the mutants. **B)** MFN2^R94Q/+^ (Het1 and Het2) spinal motor neurons have a lower number of mitochondria per micron in distal regions of the axon compared to wild-type (Ctrl1 and Ctrl2) counterparts. **C)** MFN2^R94Q/+^ (Het1 and Het2) motor neurons have an increased inter-mitochondrial distance in distal regions of the axon compared to wild-type (Ctrl1 and Ctrl2) counterparts. **D)** MFN2^R94Q/+^ (Het1 and Het2) show significantly decreased aspect ratio of mitochondrial in distal parts of motor neurons. (A-C) Data shown are the mean ± SD of 3 biological repeats. Each repeat contained N=10 cells analysed. Violin plots shown in E represent data for individual mitochondria from at least 18 neurons for each genotype.

### HDAC6 inhibition improves mitochondrial trafficking in MFN2^R94Q/+^ hESC-derived spinal motor neurons

Several studies pointed to HDAC6 inhibition as a promising approach to alleviating mitochondrial trafficking defects (d’Ydewalle et al., 2011; Dompierre et al., 2007; Guo et al., 2017). However, the efficacy of HDAC6 inhibition in alleviating trafficking impairment underpinned by MFN2^R94Q^ remains unknown. Here, we utilised our hESC-derived MFN2^R94Q/+^ spinal motor neurons to test the effect of a selective HDAC6 inhibitor, ACY-738 (Jochems et al., 2015), on mitochondrial trafficking within axons.

To determine the optimal concentration of ACY-738 for mitochondrial trafficking experiments, we first performed Western blot analysis of acetylated tubulin at increasing doses of ACY-738 (**Supplementary Figure 2A**). We chose 100nM ACY-738 for further experiments, as this dose increased acetylated tubulin in hESC-derived motor neurons (**Supplementary Figure 2B**). Next, we transfected MFN2^R94Q/+^ Het1 and Het2 hESC-derived motor neurons and their MFN2^+/+^ Ctrl1 and MFN2^+/+^ Ctrl2 counterparts with Mito-DsRed2 plasmid to label mitochondria and eGFP plasmid to label neurons. Transfected cells were treated for 24h with 100nM ACY-738 before imaging of DsRed2-labelled mitochondria by time-lapse microscopy. Strikingly, time-lapse analysis and tracking of mitochondria revealed an increase in the percentage of motile mitochondria in ACY-738-treated MFN2^R94Q/+^ cells (**Figure 5A**). The effect of ACY-738 treatment was apparent for both anterograde (**Figure 5B**) and retrograde (**Figure 5C**) movement of mitochondria within MFN2^R94Q/+^ axons. Furthermore, the aspect ratio MFN2^R94Q/+^ motor neurons in the more distal parts of the axons was significantly increased when treated with ACY-738 compared with their untreated counterparts **(Figure 5D)**, indicating that increased fusion has occurred, thus increasing mitochondrial size. Overall, this data shows that inhibition of HDAC6 using ACY-738 rescues defects in mitochondrial transport in MFN2^R94Q/+^ hESC-derived spinal motor neurons, and restores mitochondrial aspect ratio in the distal axon.

**Figure 5.**
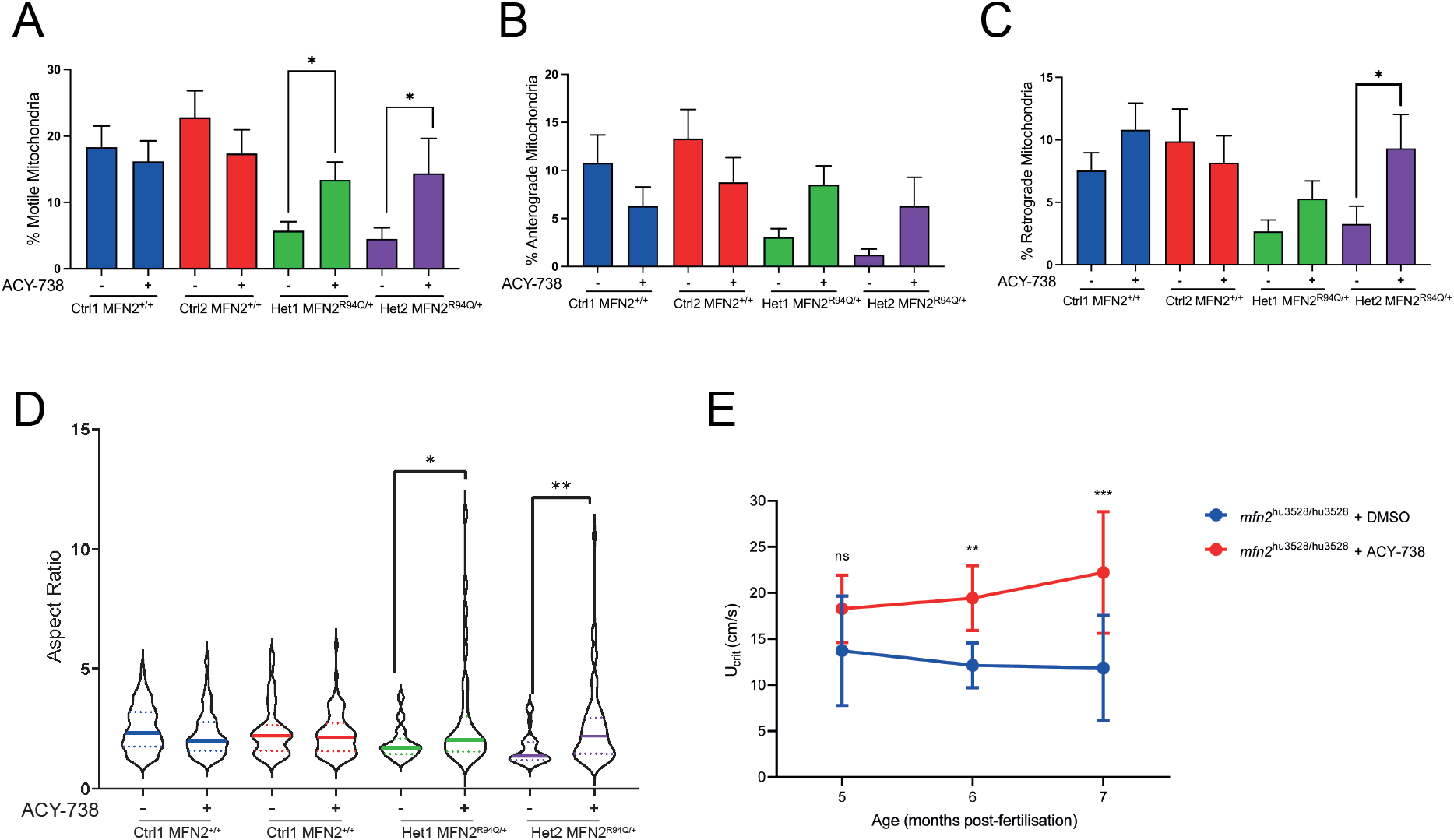
ACY-738 reverses trafficking and morphological defects in MFN2^R94Q/+^ motor neurons, as well as motor defects in MFN2 mutant zebrafish. **A)** The percentage of motile mitochondria is increased in MFN2^R94Q/+^ (Het1 and Het2) neurons upon ACY-738 treatment. **B)** The percentage of motile mitochondria in control and mutant lines moving in anterograde direction with and without ACY-738 treatment. **C)** The percentage of motile mitochondria in control and mutant lines moving in retrograde direction ACY-738 treatment. Data in A-C are the mean ±SEM. At least 15 neurons for each condition/genotype were analysed. *p<0.05, Kolmogorov-Smirnov test. **D)** The aspect ratio of mitochondria in distal parts of the axon are significantly increased upon treatment with ACY-738. Violin plots contain individual data for mitochondrion from at least 15 neurons for each condition. *p<0.05; **p<0.01, Kolmogorov-Smirnov test. **E)** ACY-738 treatment improves swimming endurance in *mfn2*^hu3528^ mutant zebrafish. The change in critical swimming velocity (U_crit_) of *mfn2*^hu3528/hu3528^ zebrafish between 5 and 7 months post-fertilisation following intermittent treatment by combined oral and immersion dosing with vehicle (1% DMSO) or ACY-738. Points and error bars represent mean and standard deviations of U_crit_ of individual fish measured at each time point (n = 8 vehicle-treated, n = 11 ACY-738-treated fish at 5 and 6 months post-fertilisation; n = 7 vehicle-treated, n = 11 ACY-738-treated fish at 7 months post-fertilisation). ns= non-significant, **p<0.01; ***p<0.001, two-way ANOVA with multiple comparisons between treatment groups at each time point.

### HDAC6 loss of function mutation transiently rescues motor defects in MFN2 mutant zebrafish

To investigate whether the potentially beneficial effect of HDAC6 inhibition in cultured MFN2^R94Q/+^ human motor neurons translates to the more complex physiological situation *in vivo* we utilised a zebrafish CMT2A model. We previously showed that *mfn2* mutant zebrafish carrying a null allele show a progressive motor phenotype associated with impaired mitochondrial transport and degeneration of the distal axon (Chapman et al., 2013). To determine whether complete loss of *hdac6* modifies this phenotype we designed experiments to generate double mutant *mfn2* and *hdac6* zebrafish and compare these with *mfn2* mutants. We previously showed that *hdac6* mutant fish have increased levels of acetylated tubulin in the majority of cells (Lysyganicz et al., 2021).

Experimental cohorts of *hdac6* ^sh400/sh400^; *mfn2*^hu3528/hu3528^, *hdac6* ^+/+^; *mfn2*^hu3528/hu3528^ as well as wild type and *hdac6* ^sh400/sh400^ controls were generated using a breeding strategy described under methods. At 3.5 months of age we compared critical swimming velocity between *hdac6* ^sh400/sh400^ and wild-type siblings. Both groups had Ucrit values >26 cm/second, suggesting there is no effect of loss of *hdac6* on swimming. At monthly intervals between 3 and 6 months old, the critical swimming velocity of *hdac6* ^sh400/sh400^; *mfn2*^hu3528/hu3528^ and *hdac6* ^+/+^; *mfn2*^hu3528/hu3528^ was determined using a swim tunnel (Chapman et al., 2013) (**Supplementary Figure 4**). Interestingly, *hdac6* ^sh400/sh400^; *mfn2*^hu3528/hu3528^ double mutants showed a significant improvement in critical swimming velocity compared to *hdac6* ^+/+^; *mfn2*^hu3528/hu3528^ zebrafish at 3 months, but this effect was lost from 4 months onwards. This suggests that loss of *hdac6* protein is beneficial in the zebrafish CMT2A model, but that the effect is transient.

### Chemical inhibition of HDAC6 rescues motor defects in MFN2 mutant zebrafish

One possible reason why the genetic ablation of *hdac6* did not restore critical swimming velocity at the later timepoints is compensation by an additional deacetylating enzyme. Such compensatory mechanisms have been described for null alleles in zebrafish and other vertebrates (El-Brolosy et al., 2019), and in keeping with this, we have observed more severe ciliary phenotypes when combining *sirt2* and *hdac6* mutant alleles in zebrafish (Lysyganicz et al., 2021). We reasoned that the chemical inhibition of HDAC6 may not invoke genetic compensation.

We performed *mfn2*^hu3528/+^ in-crosses and randomly separated the offspring into two treatment groups which were housed in separate tanks that were not connected to the recirculating water system in the aquarium. From 8 days post fertilisation, the control group were accommodated in tank water containing 2% DMSO and the treatment group were housed similarly, but the tank water also contained 300nM ACY-738. To minimise the possibility of off-target and toxic effects, an intermittent dosing strategy was used, alternating ACY-738/vehicle-treated water with untreated water on a bi-weekly basis. Despite these modifications, we noticed that the zebrafish housed in these tanks developed at a much slower rate than those maintained on the recirculating aquarium system.

Oral dosing of vehicle or ACY-738 was commenced in the diet from 80 days post fertilisation, and was also administered on an intermittent basis, alternating with untreated food on the same bi-weekly schedule as the immersion treatment. The vehicle treatment group were fed 16mg of gelly belly diet twice per day, and the treatment group were fed 16mg of gelly belly diet containing 0.058% (w/w) ACY-738 twice per day. We investigated the critical swimming velocity of ACY-738- and vehicle-treated fish at monthly intervals between 5- and 7-months post fertilisation, and Ucrit was calculated for both treatment groups at each time-point. From 6 months, the Ucrit of ACY-738-treated fish was significantly increased compared to the *mfn2*^hu3528/hu3528^ mutants, indicating that ACY-738 treatment has improved the defective swimming phenotype (**Figure 6E**). At 7 months, the experiment was terminated on welfare grounds, as the vehicle treatment animals were showing signs of distress. Together, these results show that ACY-738 treatment restores defective swimming in a zebrafish CMT2A model and that the beneficial effect is more long-lasting than we observed in a hdac6 null background.

## Discussion

To date there are no approved treatments for CMT2A and further therapeutic screening is required to identify targets which may provide alleviation of disease symptoms for patients. Reliable disease models that recapitulate key features of the disease phenotype are critical for deciphering disease pathophysiology and uncovering therapeutic options. In this study, we capitalised on hESC technology to establish an isogenic human cell-based model of CMT2A. To this end, we combined the CRISPR/Cas9 genome editing and directed differentiation of hESCs to derive isogenic spinal motor neurons harbouring one of the most prevalent and phenotypically severe CMT2A-causing mutations, MFN2^R94Q/+^. We then utilised this cell-based model to analyse CMT2A disease pathogenesis and test a pharmacological approach to alleviating the disease phenotype.

Recently, two studies have described hiPSC-based models of CMT2A (Saporta et al., 2015; Van Lent et al., 2021). In contrast to these studies, which examined patient-derived hiPSCs, our approach entailed editing MFN2^+/+^ hESCs to generate MFN2^R94Q/+^ mutant cells. There are advantages and disadvantages to both hiPSC and hESC-based approaches for disease modelling. For example, hiPSC-based approach utilises patient-specific cells, thus reflecting the exact genotype of the patient, which can be advantageous particularly in diseases where genetic modifiers are thought to play a role in a disease’s phenotype and its penetrance. However, hiPSC models are known to be highly variable due to a range of factors, including differences between donors, reprogramming techniques and genetic changes acquired through reprogramming or culturing of hiPSCs (Volpato and Webber, 2020). The use of skin biopsies from patients may also add to this complexity. Specifically, skin fibroblasts contain a range of somatic UV-related mutations that can be captured in hiPSCs during reprogramming (Rouhani, 2021) and these may affect the disease phenotype of the hiPSCs or their differentiated derivatives, despite being unrelated to the genotype of interest. In contrast, the use of hESCs combined with specific targeting of the genome allows the creation of very precise genome alterations with less likelihood of the introduction of additional unwanted mutations. Thus, here we utilised relatively early-passage hESCs (around 30 passages from an established hESC line in culture), and we generated independent MFN2^R94Q/+^ clones. After screening for potential off-target effects and karyotype, we proceeded with two clones which we then monitored regularly (every 10 passages) for the presence of common culture-acquired karyotypic abnormalities. It is important to note that the mutant clones were not whole-genome sequenced nor assessed for potential epigenetic changes. Nonetheless, by using two clones generated in two independent CRISPR/Cas9 targeting experiments, we strived to minimise the risks of the culture-acquired (epi)genetic aberrations affecting the phenotype. Importantly, hESC-based approach utilised here allows generation of disease models without the need for a patient sample, which may be particularly beneficial in the case of modelling rare genotypes or generating an allelic series on the same genetic background. Moreover, the recapitulation of the disease phenotype upon introduction of the disease-associated mutation into wild-type hESCs serves as a good validation of the disease-causing role of a particular variant and can be used to assess variants of unknown significance. Ultimately, hESC-based approach utilised here is complementary to hiPSC-based approach and provides a powerful platform for disease modelling.

Pleiotropic cellular functions of MFN2 have made it difficult to discern the cellular mechanisms through which mutations in MFN2 lead to CMT2A pathogenesis. Using the time-lapse tracking of labelled mitochondria, we showed that MFN2^R94Q/+^ human motor neurons have impaired mitochondrial axonal trafficking as they contained less motile mitochondria compared to wild-type cells. We also detected significantly smaller mitochondria in distal parts of the mutant axons. Van Lent et al (2021) have also recently described mitochondrial trafficking defects and altered mitochondrial aspect ratio in a CMT2A MFN2^R94Q/+^ hiPSC-based model. Thus, data available so far supports the hypothesis that mitochondrial trafficking defects underlie the pathology of CMT2A. Further work in this area is needed to address the molecular and cellular mechanisms implicated. For example, our findings that the frequency of mitochondrial pausing is increased in mutant neurons may suggest issues with MFN^R94Q^ protein interaction with the Miro/Milton complex (Misko et al., 2010). Specifically, although previous work has shown via co-immunoprecipitation that MFN2 containing R94Q mutation was still able to bind Miro/Milton and did not disrupt Miro/Milton binding motor proteins (Misko et al., 2010), it is conceivable that the strength of this interaction is compromised upon R94Q substitution in MFN2, making it easier for mitochondrial cargo to be dropped from Miro, therefore halting mitochondrial transport.

Finding a drug that rescues an aberrant motor neuron phenotype could provide a platform for developing pharmacological treatments for CMT2A patients. Based on our identification of aberrant mitochondrial trafficking as a key phenotype in CMT2A human motor neurons, we focused on finding a drug that could revert this aberration. Mitochondrial axonal transport can be increased by treatment of neurons with HDAC6 inhibitors, which are thought to mediate their effects on mitochondrial trafficking through increasing acetylation of tubulin (Benoy et al., 2018; Dompierre et al., 2007; Guo et al., 2017; Mo et al., 2018). Using an HDAC6 inhibitor AC-Y738, we showed a partial reversion of the diseased phenotype, as ACY-738-treated MFN2^R94Q/+^ cells exhibited increased percentage of motile mitochondria compared to non-treated MFN2^R94Q/+^ motor neurons. To determine whether this effect on axonal transport was clinically relevant, we utilised a zebrafish CMT2A model with loss of function of mfn2. ACY-738 also alleviated phenotypic defects in the zebrafish model of CMT2A, further supporting the idea that HDAC6 inhibition is a promising route to developing a CMT2A therapy. The use of hESCs and zebrafish in this study demonstrates the value of combining *in vitro* and *in vivo* model systems to corroborate the efficacy of particular treatments.

In conclusion, our data demonstrate that MFN2^R94Q/+^ hESC-derived motor neurons exhibit mitochondrial transport defects that can be alleviated by pharmacological manipulation. The model developed in our study provides an excellent platform for identifying drugs that are likely to be clinically relevant, as we demonstrated efficacy in human cells and also in a vertebrate model of CMT2A.

## Materials and Methods

### Human pluripotent stem cell maintenance

HESC line used in this study was MShef11 human embryonic stem cell line (Thompson et al., 2020). A working bank of MShef11 was made from cells around 30 passages from the original derivation. In this study, cells were defrosted from a working bank and were grown for no longer than 10 passages from defrosting. Cells were grown in Essential 8 (made in-house based on (Chen et al., 2011)) on Vitronectin (Life Technologies), and were maintained at 37°C under a humidified atmosphere of 5% CO_2_ in air. Monitoring for culture-acquired karyotypic abnormalities was performed using G-banding after thawing the cells. G-banding was performed as previously described (Baker et al., 2016; Laing et al., 2019) with 20 metaphases analysed for each sample. After 10 passages in culture, cells were either karyotyped or checked by qPCR for presence of the most common culture-acquired karyotypic abnormalities (1q, 12p, 17q and 20q), as previously described (Baker et al., 2016; Laing et al., 2019).

### Zebrafish

All zebrafish were maintained in the University of Sheffield Bateson Centre Aquaria at 28°C. Experiments were conducted according to UK law under project license 40/8309 granted to AJG, and were subject to a local ethical review process. *mfn2*^hu3528/+^ and *hdac6*^sh400/+^ zebrafish have been previously described (Chapman et al., 2013; Lysyganicz et al., 2021). *mfn2* and *hdac6* are both located on chromosome 8 in zebrafish, therefore offspring of *mfn2*^hu3528/+^ ; *hdac6*^sh400/+^ in-crosses did not show Mendelian ratios. In order to generate experimental cohorts, *mfn2*^hu3528/+^ zebrafish were crossed with *hdac6*^sh400/sh400^, and *mfn2*^hu3528/+^ ; *hdac6*^sh400/-^ offspring were identified by genotyping. These fish were then crossed with *hdac6*^sh400/-^ to generate *hdac6*^+/+^ ; *mfn2*^hu3528/+^ and *hdac6*^sh400/sh400^ ; *mfn2*^hu3528/+^ progeny. For experimental cohorts, *hdac6*^+/+^ ; *mfn2*^hu3528/hu3528^ zebrafish were derived from *hdac6*^+/+^ ; *mfn2*^hu3528/+^ in-crosses, and *hdac6* ^sh400/sh400^; *mfn2*^hu3528/hu3528^ double mutants were derived from *mfn2*^hu3528/+^ ; *hdac6*^sh400/sh400^ in-crosses (thus the experimental fish are not siblings, but share common grandparents). Wild type and *hdac6*^sh400/sh400^ single mutant zebrafish from these crosses were also utilised as controls.

For drug dosing studies zebrafish were housed in 3.5 L tanks from 5 days to 6 months old at a maximum stocking density of 10 fish per litre. Beginning at 8 days old, ACY-738 / 2% DMSO (treatment) or 2% DMSO (vehicle) were given intermittently. Treatment or vehicle were applied for 3.5 days and then withdrawn for 3.5 days, to reduce the chances of toxicity. For oral dosing ACY-738 was also formulated in Gelly Belly™ food mix (Florida Aqua Farms) according to a previously described method (Sciarra J, 2014). In preliminary studies we established that a 3.5 L tank of zebrafish, at a stocking density of 10 fish per litre, would eat at most 16mg of gelly belly diet twice per day. Thus ACY-738 was formulated at a dose of 0.058% (w/w) in gelly belly and 16mg of this was fed to each 3.5 L tank twice per day. The vehicle group received gelly belly diet without ACY-738. Swimming endurance was measured using a custom-built swim tunnel apparatus and critical swimming speed (Ucrit) was calculated as previously described (Chapman et al., 2013).

### CRISPR/Cas9 genome editing

Guides and repair template were designed to exon 3 of *MFN2* using Horizon CRISPR design tool (https://horizondiscovery.com/products/tools/CRISPR-Targeted-Gene-Designer) (cRNA sequence: TCAGTGAGGTGCTGGCTCGG Repair Template: AAGTGAGAGGCATCAGTGAGGTGCTGGCTCAGAGGCACATGAAAGTGGCTTTTTTTGGCCG). cRNA and ALT-R® CRISPR-Cas9 tracrRNA (IDT, 1072532) were mixed in equimolar concentrations and heated to 95°C for 5 minutes. The ALT-R® S.p. MShef11 cells were transfected with HiFi Cas9 Nuclease V3 (IDT), cRNA:tracrRNA complex and repair template using a DigitalBio Microporator (ThermoFisher) with the following settings: 1400V, 20ms, 1 pulse. Cells were grown in E8 supplemented with 10μM Y-27632 for 48 h. Cells were then single-cell sorted using the FACS Jazz (BD Biosciences) into 96 well plates onto a feeder layer of mouse embryonic fibroblasts. After approximately 14 days of growth, a third of the well contents was transferred to the new culture vessel and transitioned to feeder-free conditions (E8 and vitronectin). The rest of the sample was taken for DNA analysis. Sanger sequencing was performed to detect the clones with the correct genome editing of MFN2. Putative off-target effects were detected in silico using IDT Guide Checker (https://eu.idtdna.com/site/order/designtool/index/CRISPR_SEQUENCE). The top 5 off-target locations (**Supplementary Table 1**) were checked by Sanger sequencing. Two heterozygous edited clones, MFN2^R94Q/+^ Het1 and MFN2^R94Q/+^ Het2, were derived in independent CRISPR/Cas9 experiments. MFN2^+/+^ Ctrl2 clone was derived from the same targeting/cloning experiment as MFN2^R94Q/+^ Het1.

### Flow cytometry analysis

Cells were harvested using TrypLE and resuspended in PBS supplemented with 10% foetal calf serum. 100,000 cells per condition were stained with primary antibody for 15 minutes at 4°C and after washing three times in PBS, they were incubated with secondary antibody for 15 minutes 4 °C in the dark. Cells were washed before analysis on BD FACS Jazz. The gate for positive cells was set using cells that were incubated with secondary antibody only. For antibodies used see **Supplementary Table 5 and 6**.

### RNA extraction, reverse transcription and quantitative real-time PCR (qPCR)

For qPCR analyses, RNA was extracted using either the Qiagen RNA easy plus (Qiagen) or the Norgen Total RNA Purification Plus Kit (Norgen). RNA was reverse transcribed using High-Capacity Reverse Transcriptase (Applied Biosystems, 4368813) according to the manufacturer’s instructions. Each 10μl qPCR reaction contained 1x TaqMan Fast Universal Master Mix (ThermoFisher, 4352042), 100nM of forward primer, 100nM of reverse primer, 100nM of probe from the universal probe library (Roche, 4683633001) and 10ng of RNA. The PCR reaction was run on QuantStudio 12K Flex Thermocycler (ThermoFisher) with the following parameters: 50°C for 2 minutes, 95°C for 10 minutes, then 40 cycles of 95°C for 15 seconds, 60°C for 1 minute. The cycle times were obtained from the QuantStudio 12K Flex Software with auto baseline settings and were then exported to Excel for copy number analysis using the 1/dCt method (1/(Ct (gene of interest) – dCt(reference gene))).

### Immunocytochemistry

Cells were fixed with 4% paraformaldehyde (PFA) for 15 minutes at room temperature. Cells were then blocked and permeabilised with 0.2% Triton X-100 (Sigma) in PBS supplemented with 10% foetal calf serum for 1hr at room temperature. Primary antibodies were incubated with samples at 4°C overnight. Secondary antibodies were incubated for 1-2 hours at room temperature in the dark. Nuclei were counterstained with Hoechst 33342 (Thermofisher) or DRAQ5 (Abcam). Stained cells were imaged using either the Incell Analyser 2000 (GE Healthcare) or LSM880 AiryScan Confocal (Zeiss). Images were analysed using Cell Profiler (Carpenter et al., 2006) using custom-made protocols. For antibodies used see **Supplementary Table 3 and 4**.

### Differentiation of hESC to spinal motor neurons

Motor neuron differentiation was optimised based on the protocols published by Maury et al. (Maury et al., 2015) and Guo et al. (Guo et al., 2017). HPSCs (3000 cells) were plated on day 0 in 96 well U-bottomed low attachment plates (Greiner) in N2B27 media (DMEM-F12 50:50 mix with Neurobasal (Gibco), N2 (Gibco) 1:100, B27 (Gibco) 1:50, Glutamax (Gibco) 1:100, Non-essential Amino Acids (Gibco) 1:100, beta-mercaptoethanol (Gibco) 1:1000) containing, 20ng/ml FGF-2 (R&D systems), 5μM Y-27632 (Geron), 0.2μM LDN193189 (Tocris), 4μM Chir99021 (Tocris), 40μM SB431542 (Tocris) and 0.05% PVA (Sigma). Plates were spun at 400g for 4 minutes. All medium changes within were replacing only 50%, media was changed every two days unless otherwise stated. At day 2, medium was changed to contain 20ng/ml FGF-2, 0.2μM LDN193189, 4μM Chir99021, 40μM SB431542, 1μM all-*trans* retinoic acid (Sigma) and 0.5μM SAG (Tocris). At day 4, medium contained 1μM all-*trans r*etinoic acid and 0.5μM SAG At day 7 medium contained 1μM all-*trans* retinoic acid, 0.5μM SAG, 10ng/ml BDNF (Peprotech) and 10ng/ml GDNF (Peprotech). At day 9, medium was changed to contain 10μM DAPT (Tocris) 10ng/ml BDNF, 10ng/ml GDNF, 1μM all-*trans* retinoic acid and 0.5μM SAG. At day 13, spheres were pooled together, dissociated using accutase and re-plated on Poly-L-Ornithine (Sigma, P4957) and geltrex-coated dishes in media containing 10μM DAPT, 10ng/ml BDNF, 10ng/ml GDNF, 0.1μM retinoic acid and 0.5μM SAG. At day 14, retinoic acid and SAG were removed from the culture media and DAPT was increased to 20μM. At day 16, 10ng/ml CNTF (Peprotech) was added to the culture media. On day 17, DAPT was removed from the media. From this point on media contained only BDNF, GDNF, CNTF at 10ng/ml. Cells were used for experiments on day 16 or 33 as stated.

### Time-lapse analysis of mitochondrial movement

Motor neurons (differentiation day 13) were seeded into geltrex-coated 35mm dishes (Ibidi, 81156) and differentiated as above. On day 33 of differentiation, neurons were transfected with p-CAG GFP (Liew et al., 2007) and DsRed2-Mito (Tanaka Bio) plasmids using a ratio of 7:3 using Lipofectamine LTX Plus (ThermoFisher, 15338100) for 5 hours, before fresh media replacement. Treatment, if applicable, was added after 1 hour. Neurons were imaged 24-48 hours post-transfection on a Zeiss LSM880 AiryScan Confocal at 37°C. For time-lapse imaging images of GFP and DsRed2 were taken every 3 seconds for 6 minutes. Kymographs were generated from time-lapse images using FIJI. Anterograde and retrograde travelling mitochondria were noted for each kymograph. Afterwards, the entire neuron was imaged using both channels Fast mode and z stacks to visualise the entire length of the neuron. Full neuron images were stitched together using FIJI image stitching. Neuron lengths were calculated using the Simple Neurite Tracer plug-in on the GFP image. The region of interest identified from the GFP image was transferred to the DsRed2-Mito image and “Analyse Particles” was run on a thresholded image to measure individual mitochondria.

### Western blotting

Cells were lysed in 1x Laemmli Buffer pre-warmed to 95°C and the total protein concentration was measure using Nanodrop (Thermofisher). Proteins (15μg/sample) were resolved by SDS-PAGE in Mini Trans-Blot Cell (Bio-Rad) with pre-stained ladder precision plus protein (BioRad). Protein was transferred to nitrocellulose membrane (BioRad) using a Mini Trans-Blot Cell (Bio-Rad). The membrane was blocked in 5% milk for 1 hour and then incubated with primary antibodies for Tubulin (CST, 1:5000) and acetylated Tubulin (Sigma, 1:5000) in PBS supplemented with 3% BSA and incubated overnight on a rolling platform at 4°C. The secondary antibody in PBS supplemented with 3% BSA (Thermofisher, Mouse 800, 1:20000 and Thermofisher, Rabbit 680, 1:20000) was incubated in the dark for 1 hour on a rolling platform at room temperature. The membrane was washed and imaged using a LiCor Odessey (LiCor).

### Electrophysiology

Whole-cell patch clamp was used to record membrane currents or membrane potentials from single cells (n = 33), at room temperature (20-25°C), using an Optopatch (Cairn Research Ltd, UK) patch clamp amplifier. The extracellular solution contained (mM): 135 NaCl, 5.8 KCl, 1.3 CaCl_2_, 0.9 MgCl_2_, 0.7 NaH_2_PO_4_, 5.6 D-glucose, 10 Hepes-NaOH, 2 sodium pyruvate. Amino acids and vitamins for Eagle’s minimal essential medium (MEM) were added from concentrates (Invitrogen, UK). The pH was adjusted to 7.5 and the osmolality was around 308 mosmol kg^-1^. Cells were viewed using an upright microscope equipped with Nomarski DIC optics (Nikon FN1, Japan) and were continuously perfused with extracellular solution. Patch electrodes were pulled from soda glass capillaries (Hilgenberg GmbH, Germany) and electrodes had resistances in extracellular solution of around 4 MΩ. The shank of the electrode was coated with surf wax (Mr Zogs Sexwax, USA) to minimise the fast electrode capacitive transients. The pipette solution contained (mM): 131 KCl, 3 MgCl_2_, 1 EGTA-KOH, 5 Na_2_ATP, 5 Hepes-KOH, 10 sodium phosphocreatine (pH 7.3, 290 mosmol kg^-1^). Voltage and current clamp protocol application and data acquisition were performed using pClamp software and a Digidata 1440A (Molecular Devices, USA). Recordings were filtered at 2.5 or 10 kHz (8-pole Bessel), sampled at 5 or 100 kHz and stored on computer for off-line analysis using Clampfit, GraphPad Prism (GraphPad Software Inc, USA) and Origin (OriginLab, USA) software. Recordings and reported currents and conductances were corrected off-line for the linear leak conductance. Membrane potentials under voltage clamp were corrected for the voltage drop across the residual series resistance (*R*_s_) at steady-state current level and for a liquid junction potential, measured between pipette and bath solutions, of –4 mV.

### Statistical analysis

Statistical analysis was performed using Graphpad Prism version 8 by statistical tests as indicated in figure legends. For all the statistical tests p<0.05 was used as the criterion for statistical significance.

## Acknowledgments

This work was partly supported by the Department of Biomedical Science studentship, the Medical Research Council (MRC) grant number MR/N009371/1 and Hereditary Neuropathy Foundation. Confocal imaging work was performed at the Wolfson Light Microscopy Facility at the School of Biological Sciences, the University of Sheffield. For the purpose of open access, the author has applied a Creative Commons Attribution (CC BY) licence to any Author Accepted Manuscript version arising.

## References

Baker, D., Hirst, A. J., Gokhale, P. J., Juarez, M. A., Williams, S., Wheeler, M., Bean, K., Allison, T. F., Moore, H. D., Andrews, P. W. et al. (2016). Detecting Genetic Mosaicism in Cultures of Human Pluripotent Stem Cells. Stem Cell Reports 7, 998–1012.

Baloh, R. H., Schmidt, R. E., Pestronk, A. and Milbrandt, J. (2007). Altered axonal mitochondrial transport in the pathogenesis of Charcot-Marie-Tooth disease from mitofusin 2 mutations. J Neurosci 27, 422–30.

Benoy, V., Van Helleputte, L., Prior, R., d’Ydewalle, C., Haeck, W., Geens, N., Scheveneels, W., Schevenels, B., Cader, M. Z., Talbot, K. et al. (2018). HDAC6 is a therapeutic target in mutant GARS-induced Charcot-Marie-Tooth disease. Brain 141, 673–687.

Carpenter, A. E., Jones, T. R., Lamprecht, M. R., Clarke, C., Kang, I. H., Friman, O., Guertin, D. A., Chang, J. H., Lindquist, R. A., Moffat, J. et al. (2006). CellProfiler: image analysis software for identifying and quantifying cell phenotypes. Genome Biol 7, R100.

Chambers, S. M., Fasano, C. A., Papapetrou, E. P., Tomishima, M., Sadelain, M. and Studer, L. (2009). Highly efficient neural conversion of human ES and iPS cells by dual inhibition of SMAD signaling. Nat Biotechnol 27, 275–80.

Chandhok, G., Lazarou, M. and Neumann, B. (2018). Structure, function, and regulation of mitofusin-2 in health and disease. Biol Rev Camb Philos Soc 93, 933–949.

Chapman, A. L., Bennett, E. J., Ramesh, T. M., De Vos, K. J. and Grierson, A. J. (2013). Axonal Transport Defects in a Mitofusin 2 Loss of Function Model of Charcot-Marie-Tooth Disease in Zebrafish. PLoS One 8, e67276.

Chen, G., Gulbranson, D. R., Hou, Z., Bolin, J. M., Ruotti, V., Probasco, M. D., Smuga-Otto, K., Howden, S. E., Diol, N. R., Propson, N. E. et al. (2011). Chemically defined conditions for human iPSC derivation and culture. Nat Methods 8, 424–9.

d’Ydewalle, C., Krishnan, J., Chiheb, D. M., Van Damme, P., Irobi, J., Kozikowski, A. P., Vanden Berghe, P., Timmerman, V., Robberecht, W. and Van Den Bosch, L. (2011). HDAC6 inhibitors reverse axonal loss in a mouse model of mutant HSPB1-induced Charcot-Marie-Tooth disease. Nat Med 17, 968–74.

Dasen, J. S., De Camilli, A., Wang, B., Tucker, P. W. and Jessell, T. M. (2008). Hox repertoires for motor neuron diversity and connectivity gated by a single accessory factor, FoxP1. Cell 134, 304–16.

Dasen, J. S. and Jessell, T. M. (2009). Hox networks and the origins of motor neuron diversity. Curr Top Dev Biol 88, 169–200.

Dasen, J. S., Liu, J. P. and Jessell, T. M. (2003). Motor neuron columnar fate imposed by sequential phases of Hox-c activity. Nature 425, 926–33.

De Vos, K. J. and Sheetz, M. P. (2007). Visualization and quantification of mitochondrial dynamics in living animal cells. Methods Cell Biol 80, 627–82.

Dompierre, J. P., Godin, J. D., Charrin, B. C., Cordelieres, F. P., King, S. J., Humbert, S. and Saudou, F. (2007). Histone deacetylase 6 inhibition compensates for the transport deficit in Huntington’s disease by increasing tubulin acetylation. J Neurosci 27, 3571–83.

Dorn, G. W., 2nd. (2020). Mitofusin 2 Dysfunction and Disease in Mice and Men. Front Physiol 11, 782.

El-Brolosy, M. A., Kontarakis, Z., Rossi, A., Kuenne, C., Gunther, S., Fukuda, N., Kikhi, K., Boezio, G. L. M., Takacs, C. M., Lai, S. L. et al. (2019). Genetic compensation triggered by mutant mRNA degradation. Nature 568, 193–197.

Fazal, R., Boeynaems, S., Swijsen, A., De Decker, M., Fumagalli, L., Moisse, M., Vanneste, J., Guo, W., Boon, R., Vercruysse, T. et al. (2021). HDAC6 inhibition restores TDP-43 pathology and axonal transport defects in human motor neurons with TARDBP mutations. EMBO J 40, e106177.

Feely, S. M., Laura, M., Siskind, C. E., Sottile, S., Davis, M., Gibbons, V. S., Reilly, M. M. and Shy, M. E. (2011). MFN2 mutations cause severe phenotypes in most patients with CMT2A. Neurology 76, 1690–6.

Gouti, M., Delile, J., Stamataki, D., Wymeersch, F. J., Huang, Y., Kleinjung, J., Wilson, V. and Briscoe, J. (2017). A Gene Regulatory Network Balances Neural and Mesoderm Specification during Vertebrate Trunk Development. Dev Cell 41, 243–261 e7.

Gouti, M., Tsakiridis, A., Wymeersch, F. J., Huang, Y., Kleinjung, J., Wilson, V. and Briscoe, J. (2014). In vitro generation of neuromesodermal progenitors reveals distinct roles for wnt signalling in the specification of spinal cord and paraxial mesoderm identity. PLoS Biol 12, e1001937.

Guo, W., Naujock, M., Fumagalli, L., Vandoorne, T., Baatsen, P., Boon, R., Ordovas, L., Patel, A., Welters, M., Vanwelden, T. et al. (2017). HDAC6 inhibition reverses axonal transport defects in motor neurons derived from FUS-ALS patients. Nat Commun 8, 861.

Jochems, J., Teegarden, S. L., Chen, Y., Boulden, J., Challis, C., Ben-Dor, G. A., Kim, S. F. and Berton, O. (2015). Enhancement of stress resilience through histone deacetylase 6-mediated regulation of glucocorticoid receptor chaperone dynamics. Biol Psychiatry 77, 345–55.

Laing, O., Halliwell, J. and Barbaric, I. (2019). Rapid PCR Assay for Detecting Common Genetic Variants Arising in Human Pluripotent Stem Cell Cultures. Curr Protoc Stem Cell Biol 49, e83.

Lee, S. K. and Pfaff, S. L. (2003). Synchronization of neurogenesis and motor neuron specification by direct coupling of bHLH and homeodomain transcription factors. Neuron 38, 731–45.

Liew, C. G., Draper, J. S., Walsh, J., Moore, H. and Andrews, P. W. (2007). Transient and stable transgene expression in human embryonic stem cells. Stem Cells 25, 1521–8.

Lysyganicz, P. K., Pooranachandran, N., Liu, X., Adamson, K. I., Zielonka, K., Elworthy, S., van Eeden, F. J., Grierson, A. J. and Malicki, J. J. (2021). Loss of Deacetylation Enzymes Hdac6 and Sirt2 Promotes Acetylation of Cytoplasmic Tubulin, but Suppresses Axonemal Acetylation in Zebrafish Cilia. Front Cell Dev Biol 9, 676214.

Mandal, A. and Drerup, C. M. (2019). Axonal Transport and Mitochondrial Function in Neurons. Front Cell Neurosci 13, 373.

Maury, Y., Come, J., Piskorowski, R. A., Salah-Mohellibi, N., Chevaleyre, V., Peschanski, M., Martinat, C. and Nedelec, S. (2015). Combinatorial analysis of developmental cues efficiently converts human pluripotent stem cells into multiple neuronal subtypes. Nat Biotechnol 33, 89–96.

Misko, A., Jiang, S., Wegorzewska, I., Milbrandt, J. and Baloh, R. H. (2010). Mitofusin 2 is necessary for transport of axonal mitochondria and interacts with the Miro/Milton complex. J Neurosci 30, 4232–40.

Mizuguchi, R., Sugimori, M., Takebayashi, H., Kosako, H., Nagao, M., Yoshida, S., Nabeshima, Y., Shimamura, K. and Nakafuku, M. (2001). Combinatorial roles of olig2 and neurogenin2 in the coordinated induction of pan-neuronal and subtype-specific properties of motoneurons. Neuron 31, 757–71.

Mo, Z., Zhao, X., Liu, H., Hu, Q., Chen, X. Q., Pham, J., Wei, N., Liu, Z., Zhou, J., Burgess, R. W. et al. (2018). Aberrant GlyRS-HDAC6 interaction linked to axonal transport deficits in Charcot-Marie-Tooth neuropathy. Nat Commun 9, 1007.

Prior, R., Van Helleputte, L., Benoy, V. and Van Den Bosch, L. (2017). Defective axonal transport: A common pathological mechanism in inherited and acquired peripheral neuropathies. Neurobiol Dis 105, 300–320.

Rayon, T., Maizels, R. J., Barrington, C. and Briscoe, J. (2021). Single-cell transcriptome profiling of the human developing spinal cord reveals a conserved genetic programme with human-specific features. Development 148.

Reilly, M. M., Murphy, S. M. and Laura, M. (2011). Charcot-Marie-Tooth disease. J Peripher Nerv Syst 16, 1–14.

Rocha, A. G., Franco, A., Krezel, A. M., Rumsey, J. M., Alberti, J. M., Knight, W. C., Biris, N., Zacharioudakis, E., Janetka, J. W., Baloh, R. H. et al. (2018). MFN2 agonists reverse mitochondrial defects in preclinical models of Charcot-Marie-Tooth disease type 2A. Science 360, 336–341.

Rossaert, E. and Van Den Bosch, L. (2020). HDAC6 inhibitors: Translating genetic and molecular insights into a therapy for axonal CMT. Brain Res 1733, 146692.

Rouhani, F. J. Z. X., Danecek, P., Dias Amarante, T., Koh, G., Wu, Q., Memari, Y., Durbin, R., Martincorena, I., Bassett, A.R., Gaffney, D., Nik-Zainal, S. (2021). Substantial somatic genomic variation and selection for BCOR mutations in human induced pluripotent stem cells. bioRxiv.

Rousso, D. L., Gaber, Z. B., Wellik, D., Morrisey, E. E. and Novitch, B. G. (2008). Coordinated actions of the forkhead protein Foxp1 and Hox proteins in the columnar organization of spinal motor neurons. Neuron 59, 226–40.

Saporta, M. A., Dang, V., Volfson, D., Zou, B., Xie, X. S., Adebola, A., Liem, R. K., Shy, M. and Dimos, J. T. (2015). Axonal Charcot-Marie-Tooth disease patient-derived motor neurons demonstrate disease-specific phenotypes including abnormal electrophysiological properties. Exp Neurol 263, 190–9.

Sciarra J T. A., Kolb A. (2014). A gelatin-based diet for oral dosing juvenile to adult zebrafish (Danio rerio).. Laboratory Animal Science Professional 4, 40–43.

Simeone, A., Acampora, D., Arcioni, L., Andrews, P. W., Boncinelli, E. and Mavilio, F. (1990). Sequential activation of HOX2 homeobox genes by retinoic acid in human embryonal carcinoma cells. Nature 346, 763–6.

Stifani, N. (2014). Motor neurons and the generation of spinal motor neuron diversity. Front Cell Neurosci 8, 293.

Stuppia, G., Rizzo, F., Riboldi, G., Del Bo, R., Nizzardo, M., Simone, C., Comi, G. P., Bresolin, N. and Corti, S. (2015). MFN2-related neuropathies: Clinical features, molecular pathogenesis and therapeutic perspectives. J Neurol Sci 356, 7–18.

Takahashi, K., Tanabe, K., Ohnuki, M., Narita, M., Ichisaka, T., Tomoda, K. and Yamanaka, S. (2007). Induction of pluripotent stem cells from adult human fibroblasts by defined factors. Cell 131, 861–72.

Thompson, O., von Meyenn, F., Hewitt, Z., Alexander, J., Wood, A., Weightman, R., Gregory, S., Krueger, F., Andrews, S., Barbaric, I. et al. (2020). Low rates of mutation in clinical grade human pluripotent stem cells under different culture conditions. Nat Commun 11, 1528.

Thomson, J. A., Itskovitz-Eldor, J., Shapiro, S. S., Waknitz, M. A., Swiergiel, J. J., Marshall, V. S. and Jones, J. M. (1998). Embryonic stem cell lines derived from human blastocysts. Science 282, 1145–7.

Van Lent, J., Verstraelen, P., Asselbergh, B., Adriaenssens, E., Mateiu, L., Verbist, C., De Winter, V., Eggermont, K., Van Den Bosch, L., De Vos, W. H. et al. (2021). Induced pluripotent stem cell-derived motor neurons of CMT type 2 patients reveal progressive mitochondrial dysfunction. Brain 144, 2471–2485.

Verhoeven, K., Claeys, K. G., Zuchner, S., Schroder, J. M., Weis, J., Ceuterick, C., Jordanova, A., Nelis, E., De Vriendt, E., Van Hul, M. et al. (2006). MFN2 mutation distribution and genotype/phenotype correlation in Charcot-Marie-Tooth type 2. Brain 129, 2093–102.

Volpato, V. and Webber, C. (2020). Addressing variability in iPSC-derived models of human disease: guidelines to promote reproducibility. Dis Model Mech 13.

Zuchner, S., Mersiyanova, I. V., Muglia, M., Bissar-Tadmouri, N., Rochelle, J., Dadali, E. L., Zappia, M., Nelis, E., Patitucci, A., Senderek, J. et al. (2004). Mutations in the mitochondrial GTPase mitofusin 2 cause Charcot-Marie-Tooth neuropathy type 2A. Nat Genet 36, 449–51.

